# Dissecting the root-fungal interface in 3D reveals spatially distinct signalling landscapes

**DOI:** 10.1101/2025.10.22.683817

**Authors:** Syona Baptista Thomas, M Sreepadmanabh, Vidha Srivastava, Abhirami Puzhakkal, Tapomoy Bhattacharjee, Amey Redkar

## Abstract

Plant rhizospheric interactions represent intricate relationships that determine plant fitness and are crucial for interrogating host-pathogen dynamics, with significant fundamental and translational implications. Most fungal-plant interactions occur in soil – a disordered and granular 3D environment – and hence remain challenging to unravel due to complex regulatory networks. Our current body of evidence characterizing these molecular dialogues largely stems from experimental systems employing soil or *in vitro* 2D flat plates, hydroponics and gnotobiotic systems. Soil itself features widely varying visco-elasto-plastic material properties, and its inherent opacity precludes direct visualization of the infection progression in complex diseases such as wilts and root rots. Here, we introduce the first such optically transparent, 3D granular growth matrix to recapitulate complex properties of the soil microenvironment, which enables direct, cellular-level visualization of the plant-fungal interface. Our mechanically tunable 3D matrices support long-term co-culture of plants and fungi with compatibility to classical molecular and physiological assays for unravelling the early signalling events and inter-kingdom crosstalk. By leveraging the optical transparency of this matrix, we track fungal development in response to host signals *ex-planta* with 3D resolution, to report pioneering evidence of hyphal reprogramming preferentially towards the root tips during the early stages of infection. Crucially, we integrate spatiotemporal transcriptomic analyses and discover distinct pathogen-host *ex-* and *in-planta* modules during early signalling, which are likely associated with biomimetic soil-like environments. Together, our findings establish an integrable and versatile 3D platform offering an unprecedented view of the pathogen infection processes, which enables fundamental discoveries into the biological regulation of growth and infection. These insights hold immense potential for advancing our understanding of host immune responses and adaptation of filamentous pathogens, as well as open avenues to decipher drought and disease-resistance mechanisms with major agricultural benefits.

## Introduction

Plants associate with a plethora of microbes across their life span, which constitutes their microbiome to modulate growth and development ^1-3^. Such microbial associations comprising of mutualistic fungi and bacteria represent ancient alliances from the Devonian period which have shaped plant terrestrialization^4^. Despite the importance of fungi in the functioning of the Earth’s ecosystem, the genetic mechanisms of fungal adaptability to occupy complex niches in the biosphere remain largely unexplored^5,6^. Fungal diseases are on the rise in plants and animals, including humans and critically affect health and nutrition. Fungal-plant interactions jeopardize crop productivity and continue to provoke 10 - 23% yield losses^7,8^. Most fungus-plant interactions occur in the soil, and they crucially affect plant health due to the interkingdom interplay that functions across the root-soil continuum^3^. Hence, the rhizosphere microbiome represents a dynamic and complex ecosystem which execute microscale biotic and abiotic processes that determine plant fitness^9^. However, the heterogenic and disordered nature of the root-soil interface, impose limitations to decipher how roots perceive and respond to environmental cues, despite being colonization hotspots^10^.

Our understanding of how 3D physical confinement in soil – that represents a visco-elasto-plastic milieu rich in organic substrate, where fungi thrive – remains limited. This is largely due to the inherent opacity and structural complexity of soil^11-13^, that precludes direct visualization of the fungal reprogramming and progression of infection in diseases such as vascular wilts and root rots caused by ascomycetous fungi such as *Fusarium oxysporum* (Fo)^14^. Previous attempts have largely focused on defining of the soil microbiota and the functional contributions of the inter-kingdom interactions to plant health^3,15^. While these approaches have advanced our understanding of the temporal dynamics and functions of rhizosphere microbes at community level^16^, they largely offer static snapshots of microbial behaviour and interactions.

Past efforts on characterizing molecular dialogues and signaling events in rhizosphere largely stems from bacterial systems employing *in vitro* 2D flat plates, hydroponics, microfluidics and natural or gnotobiotic systems^17,18^. Also, transparent soil-like systems have enabled visualization of microbial interactions on roots, in soil-like biomimetic platforms^19-23^. Most of these previous studies have uncover prokaryotic interactions, at a system level understanding through bulk colonization. Using cryolite-based granular matrices mimicking soil-like microenvironment, soil texture has recently been shown to impact bacterial motility and migration towards their establishment on plant roots^24^. However, none of these platforms support optimal conditions for co-culturing host and microbial partners, to understand the real-time interaction dynamics, providing physiological and transcriptional snapshots in 3D across scales, along the root axis.

Physical constraints imposed by the soil granules are likely to modulate fungal growth and morphogenesis which represents varied unicellular to filamentous forms during their invasive growth. As fungi are highly adaptive to anthropogenic environmental changes, it is conceivable that mechanical and physical constraints may probe their cellular signalling and have an influence on their host interaction outcomes^25^. Recent studies have shown the substantial physical forces which fungi exert to fulfil their lifestyle of invasive growth, primarily after a physical contact on the aerial plant tissues^26-28^. Even though the role of fungi has been widely studied primarily as decomposers and mediators of nutrient and water exchanges with their hosts in soil, the underlying regulatory networks of fungal adaptation *ex-planta*, remains largely unexplored. Recent report has shed some light on how symbiotic fungi modulate their anatomical architecture and transport across hyphal networks to meet the trade demands^29^. Also, modeling has shown the biophysical metrics of the tubular nature of fungal growth during hyphal branching and nutrient growth in 3D^30^. Roots actively respond to adaptative growth responses and monitor tissue integrity by sensing gas diffusion in the environment^31,32^. Yet, the molecular mechanisms governing hyphal reprogramming under physical confinement, to sense their host before the physical contact and spatiotemporal immunity modules from host that counteract such mechanisms remain poorly understood.

To discover such mechanisms, we pioneered an optically transparent, 3D granular growth matrix to recapitulate complex properties of the soil microenvironment, which enables direct, cellular-level visualization of the plant-fungal interface. Our mechanically tunable 3D matrices support long-term monitoring of a pathosystem showing systemic infection causing vascular wilt caused by Fo^33^ and reveal spatial patterns of hyphal reprogramming and distinct pathogen-host *ex-* and *in-planta* modules, defining transcriptional states along the roots, that potentially defines the boundaries for rhizosphere interactions. Together, our findings provide an unprecedented view of the soil-borne fungal infection, going beyond traditional setups, by integrating engineered 3D models with biophysical frameworks and molecular techniques to decipher emerging fungal diseases.

## Results

### A granular viscoelastic 3D platform for interrogating plant growth under soil-like mechanical regimes

The root-soil interface microenvironment harbor complex multi-organismic interactions across microbial communities and this interkingdom balance is crucial for determining plant health (**Fig. 1a**). However, opacity of the natural soil limits our current capability of directly visualizing interactions between plant root and fungal cells inside a granular 3D environment. To engineer a transparent 3D growth media which captures the porous and granular nature of soil, we manufactured jammed packings of hydrogel granules which exhibit viscoelastic properties and a soft solid-like behaviour. By leveraging a flash solidification-based synthesis approach, we generated micron-scale solid granules of agarose—a biocompatible and chemically inert polymeric hydrogel (**Fig. 1b-c**). Given that the internal mesh sizes of polymeric agarose are on the order of ∼100 nm, these granules allow unimpeded, uniform diffusion of small molecules such as nutrients and oxygen throughout the bulk system. Effectively, our platform presents a granular and porous 3D growth matrix resembling mechanical regimes likely to be encountered in soil. Interestingly, a large internal mesh size results in a minimal scattering rendering an optically transparent matrix.

**Figure 1.**
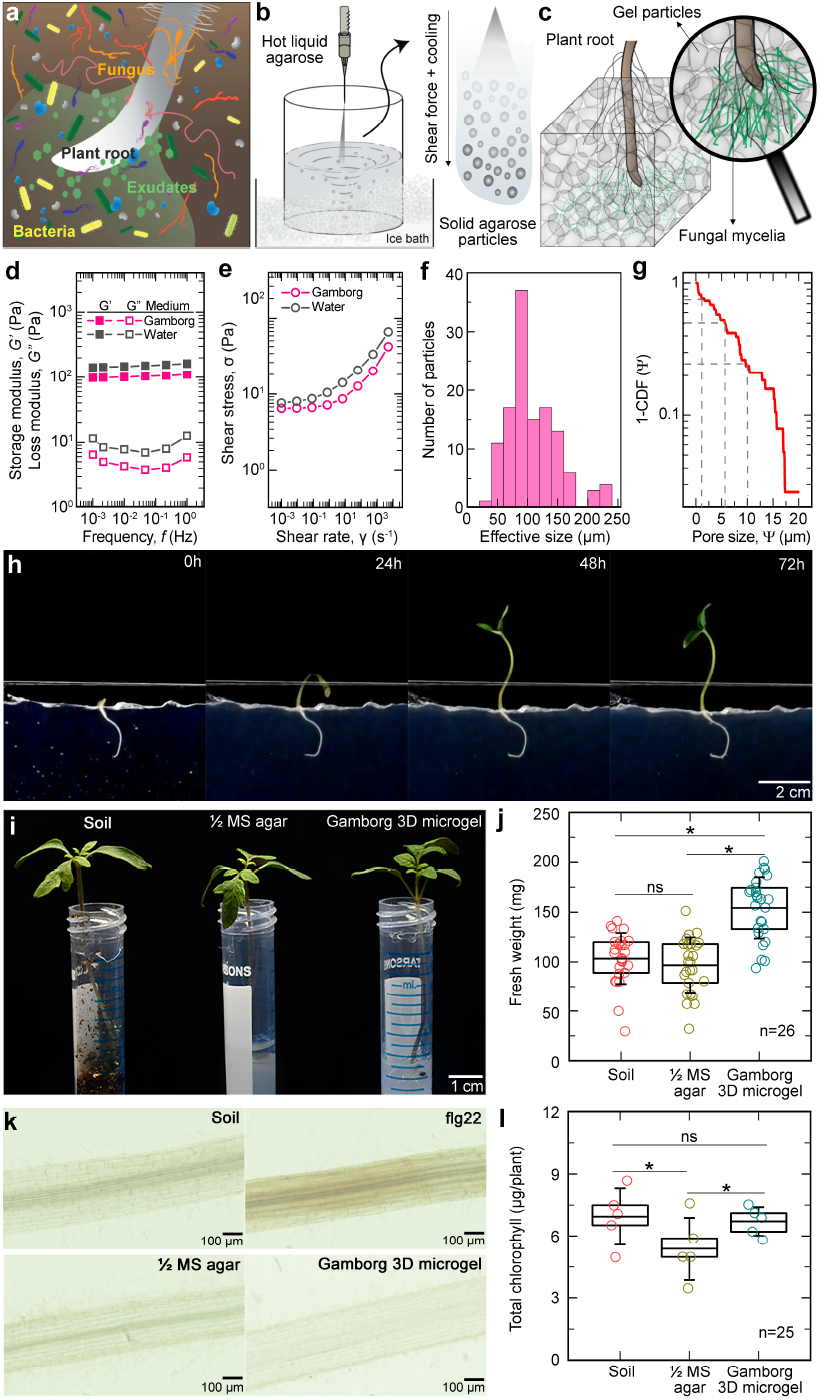
An optically transparent 3D soil-like growth matrix with tunable viscoelastic properties for plant growth. **a**, Schematic representation of the root-soil interface microenvironment, depicting the complex interactions of microbial communities with plant roots in presence of other chemical components that determine the interaction outcomes. **b**, Engineering optically transparent 3D soil-like viscoelastic growth media using flash-solidification. Molten agarose droplets are rapidly injected into ice-cold Gamborg nutrient media maintained at a high Reynolds number by vigorous stirring. The combination of shear force and cooling induces rapid gelation of these droplets into solidified granules, which are subsequently centrifuged to obtain a dense packing. **c**, jammed packings generated by flash-solidified agarose granules form a 3D granular matrix, which serves as a versatile platform for interrogating plant and fungal growth. This 3D growth media provides an internally disordered inter-particle pore-space geometry similar to a granular soil-like milieu, which supports both root growth as well as fungal hyphae and mycelia development. **d-g**, Rheological characterization of 3D growth media prepared using either Gamborg nutrient broth or water as the continuous phase. **d**, Oscillatory shear rheology-based quantification of the viscoelastic properties of the 3D growth media reveals a soft solid-like nature, with the elastic storage modulus (G’) being greater than the viscous loss modulus (G’’). Importantly, these properties are conserved across different frequencies of shear, implying that the material properties are time-invariant. **e**, Recording the stress response to varying rates of unidirectional shear reveals the yield-stress nature of the 3D growth media, which transitions from a fluid-like behavior under high shear regimes to a solid-like behavior under low shear regimes. **f**, Morphological analysis of the microgel particles indicates a polydisperse size distribution. **g**, Pore size characterization of the 3D growth matrix, measured by tracking thermal diffusion of 200 nm tracer beads within the inter-particle pore spaces, represented as a complementary cumulative distribution function (1-CDF). **h**, Representative image showing time-lapse imaging capturing real-time growth of tomato plants, from seed emergence to formation of a tomato seedling in Gamborg-based 3D growth media. **i**, Representative images showing growth of tomato plants in different substrates – soil, ½ MS and Gamborg 3D microgel – 14 days after sowing (das). **j**, DAB (3,3′-diaminobenzidine) staining of tomato roots grown in soil, ½ MS and Gamborg 3D microgel to measure their stress levels by ROS. flg22 treated root is used as a positive control. **k**, Fresh weight (mg) of the tomato roots grown in soil, 1/2 MS gel or Gamborg 3D microgel. Fresh weight of 26 individual plants measured two weeks after sowing. Box plots show the mean and interquartile range (IQR; 25th to 75th percentiles). Error bars represent the standard error mean (SEM). Statistical significance has been assessed using Student’s t-tests. p < 0.05 (*), ns = not significant. **l**, Chlorophyll content measured using methanol extraction method. Chlorophyll is quantified as µg, normalised to number of plants per sample. n=5, 5 plants per/replicate. Results are represented from five replicates. Box plots show the mean and interquartile range (IQR; 25th to 75th percentiles). Error bars represent the standard error mean (SEM). Statistical significance has been assessed using Student’s t-tests. p < 0.05 (*), ns = not significant.

To verify if the jammed packings of agarose granules act as an elastic solid—a necessary determinant of the mechanical robustness of soil— we characterize the rheological properties of these matrices. Here, we record the complex shear modulus while applying a small amplitude (1%) of oscillatory shear, which shows the storage modulus, G’ (indicative of elastic solid-like behaviour) to be consistently greater than the loss modulus, G” (indicative of viscous fluid-like behavior) (**Fig. 1d, SI Fig. 1a**). Importantly, we find that the rheological properties of this system are time-independent, remaining invariant across different shearing frequencies. Furthermore, to test whether plant roots can navigate through these porous matrices without building local stress responses, we investigate how these jammed microgel systems respond to large amplitude unidirectional shear. By subjecting the 3D growth media to varying unidirectional shear rates and recording the consequent stress responses, we find that the material exhibits yield stress behaviour by reversibly transitioning between a solid-like nature under low shear regimes and a fluid-like nature under high shear regimes (**Fig. 1e, SI Fig. 1b and 1c**). Further, we characterize the internal pore space geometry of the 3D growth media by quantifying both the individual microparticle morphologies (**Fig. 1f and SI Fig. 1d-h**) as well as the inter-particle pore size distributions. We find that jammed packings of these hydrogel granules give rise to heterogeneously distributed inter-particle pore spaces spanning several micrometers - precisely, ranging from 1 to 20 microns, with the mean pore size being approximately 5 microns **(Fig. 1g)**. Together, these results establish our 3D growth media as a granular, porous, solid-like viscoelastic material which can reversibly undergo shear-induced fluidization without a loss in its material properties.

We next assess the suitability of our engineered 3D growth media as a platform supporting plant growth. Leveraging the optical transparency of this matrix, we accomplish the real-time visualization of germination and subsequent growth of tomato seedlings across a 72 hrs. timeline from a fully embedded seed, within a soil-like granular mechanical milieu **(Fig. 1h)**. To benchmark our material against conventional experimental standards, we next performed a comparative analysis of the 3D media grown tomato seedlings with plants grown in natural soil, ½ MS gel, and the 3D growth media **(Fig. 1i)**. Phenotypic analysis of the seedling development across these tested substrates reveals no detectable effect on vegetative growth, and comparable fresh weight biomass **(Fig. 1j)**. Also, measurement of the stress-indicative reactive oxygen species (ROS) by 3,3′-diaminobenzidine (DAB) staining-a chromogenic substrate to visualize peroxidase enzymes, as well as measurement of chlorophyll content show comparable readouts demonstrating the 3D growth media as a highly suitable platform for supporting plant growth **(Fig. 1k-l)**. Taken together, these observations establish our 3D growth media as a state-of-the-art experimental system for interrogating plant biology across soil-like granular viscoelastic regimes.

### Physical confinement within soil-like 3D growth media alters the *F. oxyporum* transcriptional landscape

A crucial determinant of plant health is the ability to resist and combat pathogenic fungal infections. The outcomes of plant-fungal interactions have significant implications towards growth success and crop yield. Consequently, the mechanisms driving these interactions have been a focal point of research. However, despite an extensive characterization of the molecular effectors regulating fungal infection and plant defenses, we lack a systematic understanding of how complex soil-like physical regimes alter these events –particularly about fungal reprogramming and directed fungal growth, in contrast to traditional axenic growth comparisons from liquid cultures. A major obstacle towards capturing these dynamics in real-time in a native confined state, is the lack of granular 3D culture platforms which can support both plant and fungal growth while enabling direct visualization. Our present innovation overcomes these limitations by enabling live microscopic visualization of *in situ* plant and directed hyphal growth of *F. oxysporum* towards the roots of the host, tomato **(SI Fig. 2a)**. Firstly, by leveraging the optical transparency of our 3D growth medium, we observe the development of fluorescent mClover3-labelled isolate of the tomato pathogenic *F. oxysporum* (f. sp. *lycopersici*-Fol) hyphae across a 24-hour time-period **(Fig. 2a)**. Development of the hyphae in presence of tomato root shows a directed growth and re-orientation towards the root tip in near field area, documented at 36 hours post inoculation (hpi) **(SI Fig. 2b)**. Next, we employ optical density-based bulk measurements of biomass production to characterize the temporal dynamics of fungal growth in gamborg based liquid and 3D growth media. Interestingly, even though the growth of hyphae in liquid gamborg and 3D growth media were comparable across the 24 hours timepoint, mycelial growth in the 3D gel exhibit a higher fluorescence at the 48-hour time point. Finally, following 72 hours of growth, both liquid and 3D gel-grown hyphae achieve comparable levels of fluorescence expression, which is indicative of active fungal proliferation **(Fig. 2b)**. Going beyond such macroscopic measurements, we also accomplish the high-resolution imaging of a single fungal spore germinating in 3D space **(Fig. 2c)**, which demonstrate a normal developmental timeline as previously observed for axenic fungal culture in liquid media. Together, these efforts demonstrate the versatility of our 3D platform, which enables the visualization and quantitation of fungal growth and development within granular soil-like mechanical regimes.

**Figure 2.**
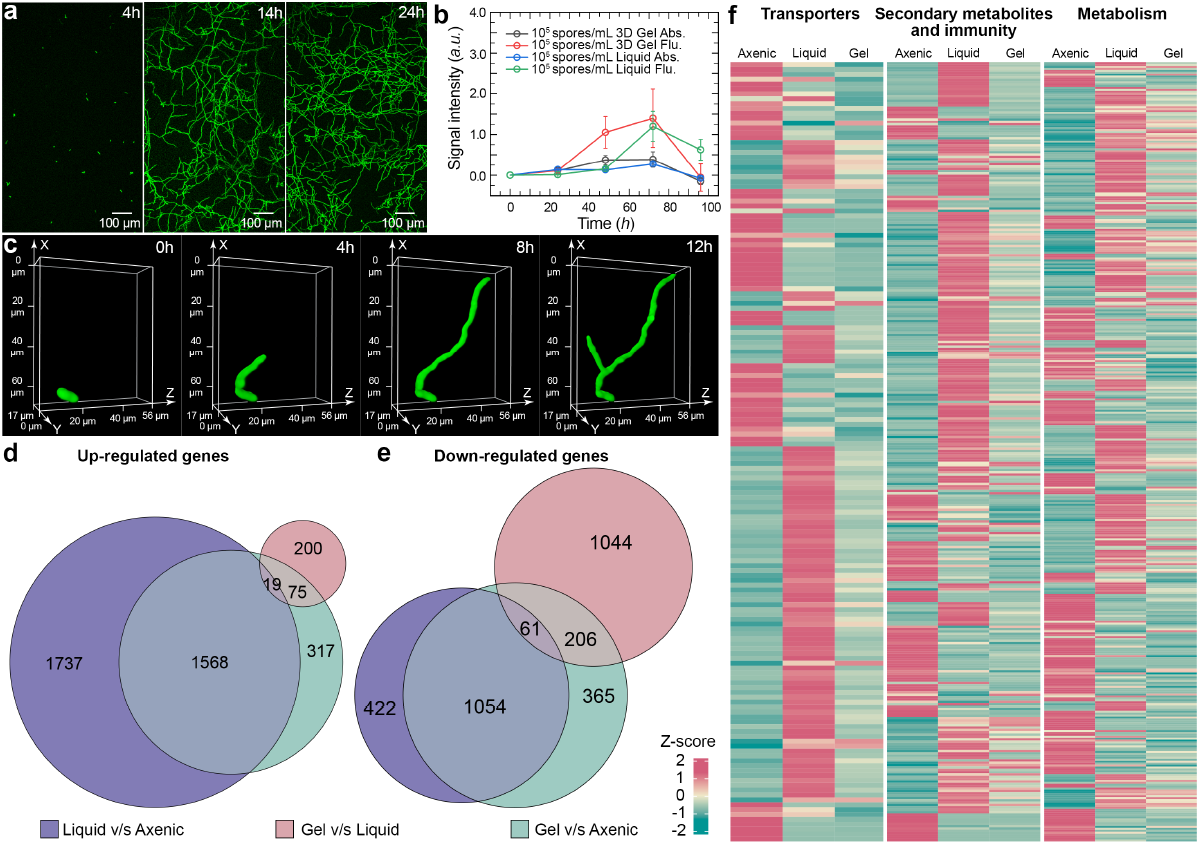
Physical confinement within soil-like 3D growth media alters the *F. oxysporum* transcriptional landscape. **a**, Fungal development starting from *F. oxysporum* f.sp. *lycopersici* (Fol)-3xmClover3 spores inoculated within the 3D growth media, imaged of over 24 hrs to track the developmental process. Scale bars, 100 µm. **b**, Growth of within liquid nutrient broth and 3D growth media, assayed using bulk fluorescence (520nm) and absorbance (600nm)-based measurements. **c**, Volumetric imaging of a single Fol-3xmClover3 spore embedded within Gamborg-based 3D growth media, demonstrating its real-time growth and development over 12 hrs. **d-e**, Euler plots representing the differentially expressed up-regulated (log2FC >2 and p-adj <0.05) (d) and down-regulated (log2FC <-2 and p-adj <0.05) genes. (e) Fol genes after DESeq2 under different growth conditions at 24 hpi. Liquid vs. axenic represents Fol grown in liquid Gamborg B5 medium, normalised to Fol grown in liquid minimal medium. Gel vs. axenic represents Fol grown in microgel jammed Gamborg B5 medium, normalised to Fol grown in liquid minimal medium. Gel vs. liquid represents Fol grown in microgel jammed Gamborg B5 medium, normalised to Fol grown in Gamborg B5 medium. All liquid cultures are shaken at 140 rpm. **f**, To better resolve the physiological basis for the differential behaviour of Fol across the three different growth conditions, we select differentially expressed protein-encoding genes in Fol, which are expressed in either liquid vs. axenic or gel vs. axenic comparisons. These are plotted with a raw z-score after running through a func-e package to perform InterPro Term Enrichment (IPR) analysis, which groups them into functional categories such as transporters, secondary metabolites and immunity, and metabolism. A marked variation in the gene expression patterns is evident for liquid vs. gel-based growth conditions. Colour scale indicates z-score.

To further investigate whether and how 3D physical confinement influences the physiological development and biological functionalities of *F. oxysporum*, we compare the transcriptomic profile of *F. oxsporum* hyphae harvested from gamborg based liquid culture and 3D growth media, following 24 hours of growth with a axenic liquid culture. Principal Component Analysis (PCA) of this data reveals that physically distinct growth conditions cluster separately, as suggested by their distinctive transcriptome profiles **(SI Fig. 3a)**. Moreover, nutritional status also reveals a major shift in transcriptional profile as seen in axenic liquid culture **(SI Fig. 3a)**. To compare growth across each of these conditions, we use a minimal nutrient media, as a reference state for axenic liquid culture, corresponding to the optimal level of gene expression during fungal development. This difference in the nutritional status also reveals a major shift in transcriptional profile as compared to the gamborg based physically distinct growth conditions **(SI Fig. 3a)**. Furthermore, we observe that growth under 3D confinement leads to distinct transcriptional landscapes in Fol as summarized with the Euler and volcano plots **(Fig. 2d-e; SI Fig. 3c-e)**. Specifically, we found 1,737 and 312 differentially expressed genes (DEGs) being upregulated in liquid and 3D growth media when compared against axenically grown Fol, respectively. However, a comparison between Fol grown in 3D growth media and liquid-grown cultures revealed only 200 upregulated genes in the 3D grown cultures compared to liquid grown cultures. Further, we find 422 and 365 genes downregulated in liquid and 3D grown gel cultures, respectively, when compared against liquid axenic (**Fig. 2d-e) (Supplementary Table 1)**. Taken together, upon comparing liquid and 3D growth media-grown cultures, we conclude that despite the overlaps observed for transcriptional responses between liquid and 3D grown gels as compared to axenic, a significant proportion of distinct DEGs also exists **(Fig. 2d-e)**. This suggests that both nutrient composition and mechanical regimes play a role in regulation of transcriptomic profiles.

**Figure 3.**
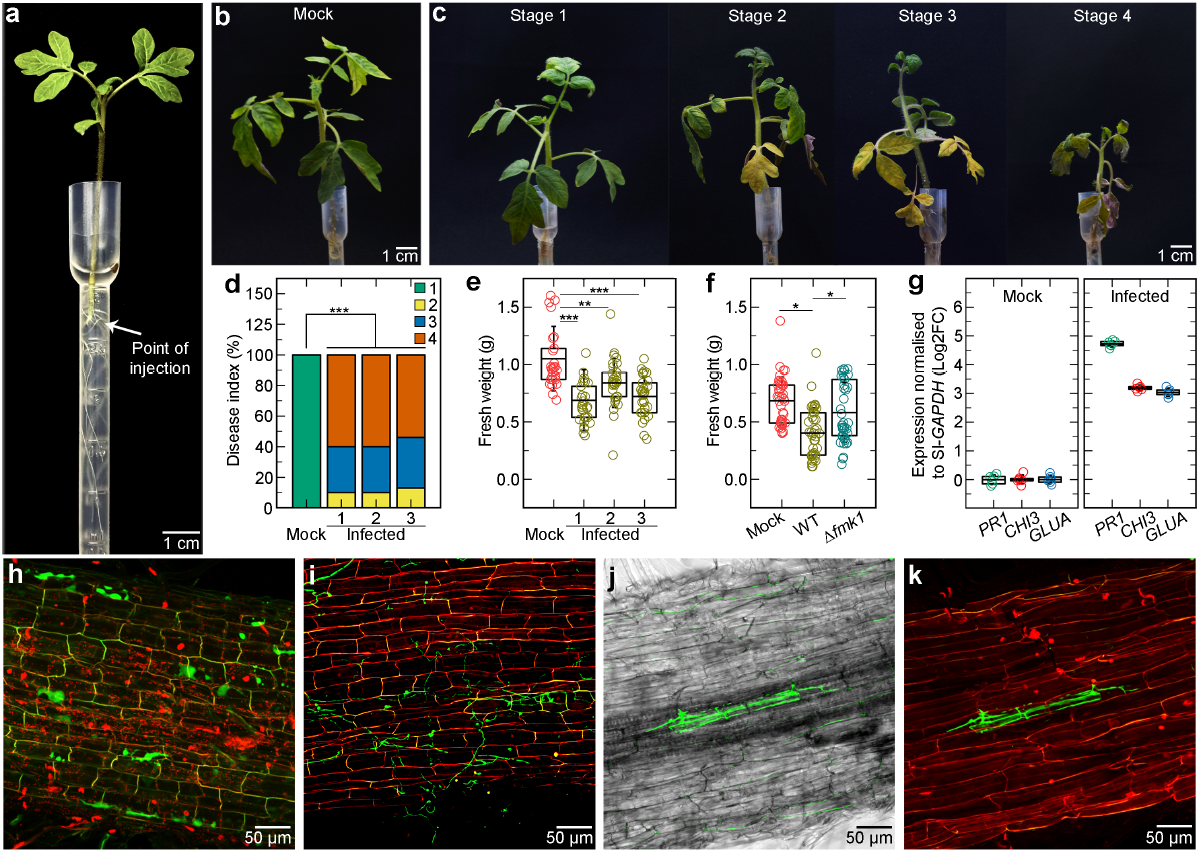
Validating the infection dynamics of vascular wilt in tomato plants using a granular 3D soil-like milieu. **a**, Representative image of a 2-week-old tomato seedling within the experimental setup for evaluation of the infection dynamics and disease symptoms in tomato infected with *F. oxysporum*. Scale bar, 1 cm. **b-c**, Scoring scheme for the progression of vascular wilt disease symptoms in tomato seedlings with the water (mock) infected tomato seedling as a reference (b) and classification of the symptoms into four stages with increasing symptom severity based on the wilting index (c). Scale bar, 1 cm. **d**, Disease symptoms of the *F. oxysporum* infected tomato seedlings scored at 35 days post infection (dpi) to evaluate the progression of the vascular-wilt disease index as a percentage of the wilted plant area. Number of independent experiments = 3; n=30 plants/treatment. *** P <0.001, versus the mock according to Chi square test. **e**, Fresh weight of tomato plants measured at 35dpi after inoculation either with water (mock) or Fol wildtype. n=30 plants/ treatment. Asterisks indicate statistical differences versus Mock (Students t test, *** P <0.001, ** P <0.01). **f**, Fresh weight of tomato plants measured at 35dpi after inoculation either with water (mock) or Fol wildtype or Δfmk1 mutant. n=40 plants/treatment. Asterisks indicate statistical differences (Students t test, * P <0.05). Data shown is a cumulative from three independent experiments. **g**, Transcript levels of tomato defence genes *pr-1, gluA* and *chi3* measured by RT-qPCR of cDNA obtained from tomato roots grown in microgel setup at 3 dpi, either with water (mock) or with the Fol wildtype (infected). Transcript levels are calculated using the ΔΔCt method, normalized to the tomato *gadph* gene and expressed relative to those of the uninoculated control (H2O). n = 5/treatment. Roots from five plants have been pooled together as one biological replicate. Experiments are performed at least two times with similar results. **h-k**, Confocal microscopy of tomato root colonization of *F. oxysporum* Fol-wildtype expressing-3xmClover3 at 3 dpi and 12 dpi, when inoculated proximal to the root surface. Fungal fluorescence (mClover3-green) is overlaid with propidium iodide staining of plant cell walls (red). Scale bars, 50 µm. Note the invasion of vasculature (xylem) at 12dpi, confirming the progressing of systemic infection in the 3D microgel setup.

However, this data does not suffice to infer whether varying stress response genes also contribute to the DEGs pool across the physically distinct growth conditions. We test this by enlisting a list of 78 stress marker genes in fungi^34^ and look for their differential patterns across our samples of physically distinct growth conditions. We find only 3 genes being upregulated and showing a similar expression pattern in both gel and liquid cultures **(SI Fig. 3b)**. Furthermore, to understand the functionalities encoded by the DEGs across confinement states, we perform an Inter-Pro Enrichment (IPR) term enrichment analysis, which identifies genes belonging to categories such as metabolism, transporters, or secondary metabolites and immunity-related gene families **(Fig. 2f, SI Fig. 3f)**. These results suggest that 3D confinement, in addition to the nutritional composition of the medium, is a major determinant of the fungal transcriptome that regulate metabolisms in response to growth state. Importantly, these findings strongly support the idea that growth under soil-like 3D regimes fundamentally differs in the transcriptional state of fungal growth as compared to homogeneous liquids by leveraging a unique, previously unidentified, set of genes. However, these differential responses are not a result of an elevated degree of stress responses but, the diverse metabolic factors that modulate growth across physically distinct conditions.

### Vascular wilt progression by *F. oxysporum* from proximity of root surface in soil-like 3D microgel

Root infection studies have classically been done by dipping roots in a spore suspension, to promote direct spore adherence to the root surface and evaluate the symptoms of a systemic disease. From the available evidence across the last decade in the soil-borne fungus such as *F. oxysporum*, it remains inconclusive whether there is any preference for the invading fungus to grow towards root tip, at the elongation zone above the root tip or across the longitudinal root surface^35-37^. Previous studies in *F. oxysporum* f.sp. *lini* report an active growth of the hyphae, within the meristematic cells of the root tip^38^. However, the frequency of such an event, where root tip encounters a fungal inoculum in soil, remains challenging to investigate. The striking differences in fungal growth and development under 3D physical confinement holds important implications for identifying the determinants of infection trigger, at a spatial resolution in a soil-like granular regimes. Understanding whether and how such mechanical properties of the soil microenvironment affect fungal sensing towards root and pathogenesis, will provide significant implications for designing resistant cultivars and developing control strategies. Motivated by this, we next investigate how vascular wilt infection by *F. oxysporum* progresses in tomato seedlings cultured using the 3D growth media, upon spot inoculation of the fungus at ∼5mm vicinity of the root elongation zone. Towards this, we grow pre-germinated seedlings in pasteur pipettes as a high-throughput mobile system, which enables us to initiate infection by directly inoculating the spore suspension in the proximity of plant roots **(Fig. 3a)**. Encouragingly, the 3D gel-grown infected plants exhibit disease symptoms showing chlorosis, wilting and ultimately result in mortality because of wilt. However, the progression of infection in such a setup is relatively slower as compared to dip infection, where spores are directly deposited on the root surface **(Fig. 3b-c)**. Importantly, at 35 days post infection (dpi) around 60% of the infected plants with wildtype Fol, show mortality as compared to the mock control. Moreover, progression of these infection symptoms are consistent across three independent replicates that evaluate pathogenicity of tomato seedlings –as indicated by wilting index and measurement of the fresh weight biomass of the individual infected plants **(Fig. 3d-e)**.

We next tested the robustness of this 3D soil-like milieu, in evaluating host plant colonization and virulence, by testing an isogenic Fol deletion mutant lacking MAPK *Fmk1* (Δ*fmk1*), which has a role in invasion growth of Fo^39^. Phenotypic analysis of the Δ*fmk1* mutant reveal significantly reduced mortality compared to the wildtype strain and an increase fresh weight biomass compared to mock, as reported previously^39^ **(Fig. 3f)**. To test whether activation of host defense occurs upon delivery of fungal inoculum proximal to the root, we measured the expression by transcript levels of known plant defense genes at 3 dpi. Upregulation of the *pr-1, gluA* and *chi3* genes encoding pathogenesis-related protein 1, basic 1,3-glucanase and acidic chitinase, respectively^40^, were 3 to 5-fold higher in tomato roots infected with the wildtype strain as compared to mock control **(Fig. 3g)**. Confocal microscopy of the infected tomato plant roots inoculated with fluorescent mClover3-labelled strain of the tomato pathogenic isolate Fol (f. sp. *lycopersici*) in the proximity of the root surface, confirmed the presence of fungal colonization in the root cortex at 3 dpi **(Fig. 3h-i)** and in xylem at 12 dpi – indicative of systemic infection **(Fig. 3j-k)**. Taken together, we conclude that our multifaceted interrogation of vascular wilt progression – combining macroscopic phenotyping, biomass quantification, host defense responses, and microscopic visualization of colonization –establish the 3D growth matrix as a versatile platform for studying fungal pathogenesis.

### Spatial transcriptomics reveals distinct fungal effector and plant defense profiles

Plant-fungal interactions at the root-soil interface **(Fig. 4a)** present a rich arena of intertwined biological regulation and transcriptional re-programming of both host and fungus, which involves plant-centric sensing of the pathogen and subsequent upregulation of immune responses, as well as remodulation of fungal states towards overcoming host defenses to establish a successful infection. Despite extensive characterizations of the underlying transcriptional signaling network driving these dynamics, we presently lack a comprehensive understanding of how distinct phenotypical manifestations trigger compatibility, through a cascade of early reprogramming events that possibility shapes these associations before a physical contact. This is in part due to the spatiotemporal complexities associated with dissecting plant-fungal interactions in rhizosphere – for instance, at what distance from the root-soil interface is microbial re-programming and differentiation for host invasion triggered? Does a host detect presence of an invading hyphae before a physical contact to prime immune responses? What are the pathogenic effectors involved in early sensing of host *ex-planta*? Does transcriptional responses across different root zones vary towards microbial invasion?

**Figure 4.**
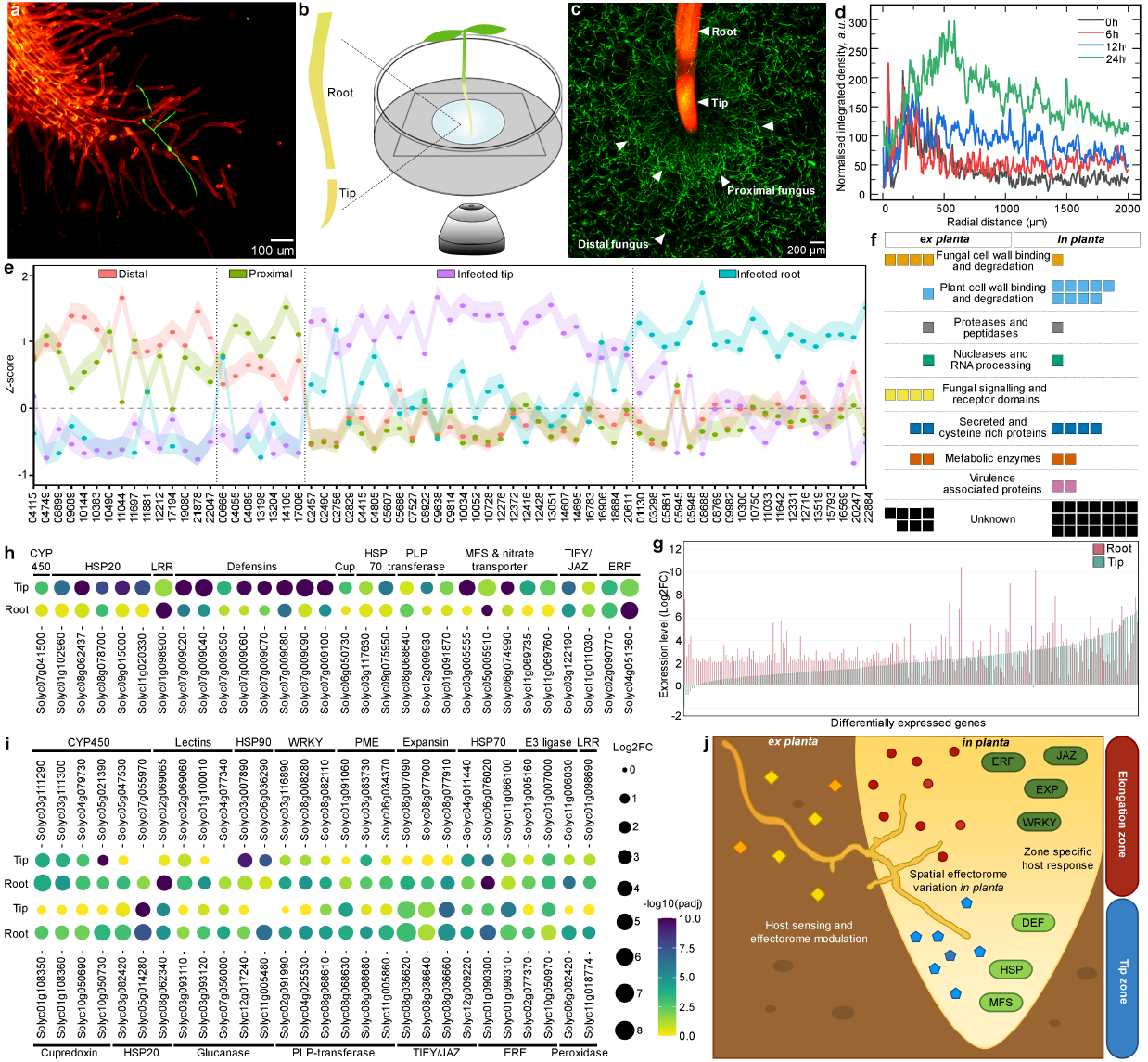
A spatiotemporal transcriptomic atlas of *F. oxysporum*-infected tomato reveals distinct *ex-* and *in-planta* signalling modules. **a**, Confocal micrograph of a tomato root showing the early interaction events with *F. oxysporum* Fol-wildtype expressing-3xmClover3 at 24 hpi in the 3D microgel setup. Fungal fluorescence (mClover3-green) overlaid with propidium iodide staining of plant cell walls (red). Scale bar, 100 µm. **b**, A schematic representation of the experimental system to capture the early interaction events in real-time at the plant fungal interface. A tomato seedling is placed centrally within a plate containing Gamborg 3D microgel co-inoculated with fungal spores. This enables parallel monitoring of the fungal infection progression towards plant roots and the consequent host responses. **c**, Confocal micrograph of the Fol-3xmClover3 germlings showing the formation of a distinct zone (indicated by arrowheads) and reorientation of the hyphal growth at 500 µm from the root surface towards the host root tip during the initial sensing phase. Scale bar, 200 µm. The sampling points for transcriptomics are for fungal samples indicated as Proximal fungus (PF), Distal fungus (DF), Root tip (RT) and Root body (RB). **d**, Quantification of normalised integrated hyphal density using 3xmClover3 fluorescence as a function of radial distance the root surface. **e**, Cleveland dot plot showing differential expression analysis of genes encoding Fol-secreted proteins plotted with a Z-score > 0.75 across samples from Proximal and Distal fungus that has re-programmed in response to host signals as well as from infected tip and root across Fol infected tomato seedling. Note a clear induction of the different subsets of the candidate effector proteins across the samples analysed. **f**, Annotation of the different subsets of the identified candidate effectors across samples ex-planta and in-planta with their IPR terms, which are grouped into functional categories. Each box represents one effector. **g**, Ranked bar plot which represents the expression of S. lycopersicum up-regulated genes (log2FC >2 and p-adj <0.05) across the spatial tissue dissection with root tip and root body, normalised to their respective control (mock tip or mock-root body). **h-i**, To understand the plant responses to the invading fungus, we perform InterPro Term Enrichment (IPR) analysis using the func-e package on genes upregulated across the root tip and root body. After removing the redundant families, the dot plot represents log2FC and the p-adj values for genes as normalised to their respective controls (mock-tip or mock-root body). (h) represents the set of genes that are highly induced in the root tip, and (i) represents the genes highly induced in the root body. **j**, Schematic representation summarising the spatiotemporal transcriptomic analyses of both the pathogen and host with distinct pathogen-host *ex-* and *in-planta* modules during early signalling, which are likely associated with biomimetic soil-like environments.

To answer such questions during the early interactions between Fol and tomato roots we designed a setup that allows us to both directly visualize these dynamics as well as retrieve samples from distinct locales **(Fig. 4b)**. Moreover, to spatially resolve host responses, we dissect the root tissue into root tip and zone of elongation **(Fig. 4b)**. Interestingly, our time-resolve fungal development in root proximity in this setup, demonstrates a marked reorientation of the fungal hyphae towards the root tip **(Fig. 4c)**. This hyphal reorientation is only seen in the presence of a plant root, indicating a reprogramming towards invasive growth **(SI Fig. 2b-c)**. Furthermore, plant root tip also causes a zonation effect on the fungal hyphae in its vicinity at ∼500 µm, as apparent from micrographs obtained 24 hours post-inoculation **(Fig. 4c-d, SI Fig. 2c)**. We note that the fungal hyphae at the elongation zone above the root tip shows invasive growth likely suggesting this zone to be the preferred for gaining an initial host entry **(Fig. 4c)**.

Next, to spatially resolve and profile the interactions between plant and fungus prior to colonization, we profile the transcriptome from fungal samples harvested near the root (proximal fungus) as well as fungal samples in the far field, away from the root surface (distal fungus). Considering that some fungal hyphae may have already entered the plant, we also compare fungal colonization profiles from both the root tip as well as the elongation zone. Principal component analysis of these samples shows that the *ex-planta* samples cluster separately as compared to the *in-planta* and to the fungus alone control without the plant host **(SI Fig. 4a)**. To identify fungal effector candidates that could shape this early dialogue, we looked for secreted DEGs in the proximal and distal fungal samples, as well as infected root and tip sections. Correlation analysis reveals that the secreted protein profiles *ex-planta*, between the proximal and distal samples are largely overlapping with each other, whereas the *in-planta* effector profiles are distinct between the tissues compared **(Fig. 4e, SI Fig. 4b-f)**. IPR term enrichment analysis shows that identified candidate effectors *ex planta*, are predicted to modulate fungal cell wall binding and degradation, signalling, and receptor domains. However, most of them account for a novel set of unannotated small secreted proteins previously unknown from *in planta* secretomes. By contrast, *in planta* secreted proteins are predicted to be involved in plant cell wall binding and include different cell wall degrading enzymes such as pectate lyase, glycoside hydrolase, gluconolactonase, peptidase, etc. Overall, our fungal proteome across samples identifies a CAZyme repertoire belonging to AA, CBM, GH and PL families **(SI Fig. 4g)**.

To identify if any of the previously identified classical effector candidates show a spatiotemporal resolution, we looked for some of the characterized fungal proteins previously known to be involved in the mechano-transduction across fungi **(Supplementary Table 4)**. Interestingly, some of the identified Fo proteins show an interesting and distinct expression pattern *ex-planta* previously not described during the colonization process. Of interest among these are the tomatinase enzyme Tom1, which is upregulated early in the *ex-planta* proximal contributing to full virulence on tomato plants^41-42^, the Lipase-Lip3 which hydrolyze the plant cuticular wax^43^, the Nitrate as well as the Nucleotide sugar transporter (NTR1 and NST6) all of which contribute as the core pathogenicity determinants^44-45^**(SI Fig. 4h)**. Together, we conclude that Fol already undergoes a host-mediated differentiation at the root-soil interface, showing a transcriptional activation by distinct secreted protein sub-sets *ex-* and *in planta*, tailored towards different root zones. This finding suggests distinct processes in pathogenic fungi facilitate sensing and exploration of host, in a disordered substrate such as soil.

The perception of host signals to modulate invasive fungal program *ex-planta* suggests that root immune responses may also be equally channelized. To test this idea, we profile the transcriptomes of infected root tip and elongation zone, relative to their respective mock control to define tomato immune transcriptional responses at the root-soil interface. The PCA shows that the root tip and elongation zone samples cluster independently from each other **(SI Fig. 4i)**, and that infected samples show a pronounced upregulation of genes relative to mock samples **(SI Fig. 4 j-l)**. A ranked bar plot of DEGs (log2FC > < 2, p-adj < 0.05) in the root tip and root body shows no correlation in expression patterns **(Fig. 4g)**. This suggests that the same set of genes are regulated differently across the two tissues, and that their overall transcriptomic profiles do not overlap. Hence, the root tip and root body activate distinct responses against Fo. IPR term enrichment analysis reveals that infected tips exhibit a strong modulation of defensins, MFSs and nitrate transporters **(Fig. 4h-i)**. Moreover, infected roots also display a strong modulation of several key defense and cell wall-related gene families. In particular, each of the WRKY transcription factors (key regulators of the plant immune system), glucanases ( which hydrolyze fungal cell wall and release elicitor fragments, reflecting an attempt from the plant to fight off the pathogen), peroxidases (indicating a localised oxidative burst at the site of infection) and, expansins (which loosen the cell wall during growth, and are possibly induced by the pathogen to weaken the cell wall and invade the plant) **(Fig. 4h-i)**. Moreover, the TIFY/JAZ gene module shows a tight regulation of its distinct members across the root tip and root body^46^ and which have previously been reported to exhibit versatile role in stress signaling. Also, *F. oxysporum* hijacks the COI-1 mediated jasmonate signaling to promote disease^47^ and hence the role of this module in early signaling is evident. Similarly, the lectin gene family also exhibits a tissue specific role and have recently shown to have immune regulatory functions in tomato^48^ **(Fig. 4h-i)**. Taken together, these results indicate that the plant responses particularly related the immune signaling are broadly activated in both the tip and elongation zone across root. However, their execution in response to fungal invasion vary in terms of expression profile representing a modular pattern **(Fig. 4j, Supplementary Table 5)**. This indicates that the distinct plant tissues likely channelize the immune responses, catering towards the tissue specialization to balance the growth-defense tradeoff. These findings reveal a foundational understanding of tissue-specific immune signalling at the root-soil interface that potentially represent a framework to understand natural infections in soil.

## Discussion

The rhizosphere represents a dynamic and complex ecosystem which executes microscale processes within a multi-kingdom microbial consortium, that critically determine plant growth and fitness^9^. The microbiome composition and the interactions within it are shaped both by the soil substrate as well as a wide range of root compounds including sugars, secondary metabolites, and in spite being an interaction hotspot, the spatio-temporal microbial dynamics in soil remain fragmentarily investigated due to its opaque nature. To address this, our work establishes a hydrogel-based granular 3D matrix as an optically transparent soil mimic, which provides a versatile platform to study plant-pathogen dynamics under controlled, yet physiologically relevant conditions^50^. Our platform offers an integrated mechanically tunable system compatible with biochemical and molecular techniques, which supports long term co-culture of plants and fungi. Here, we demonstrate its versatile capabilities for characterizing plant and fungal growth, interrogating vascular-wilt disease progression in real-time, as well as accomplishing both a system and cellular-level visualization of infection dynamics at the plant-fungal interface.

Previous attempts have largely defined the microbiome composition, revealing the complex nature of these microcosms by high-throughput sequencing techniques^51^. However, these efforts have profiled microbiome dynamics as a snapshot at the community level, rather than elucidating real-time mechanistic components that contribute towards determining inter-organismic interaction outcomes^52^. Moreover, biophysical studies have shown that 3D confinement significant impacts microbial activity, either by determining the colony structure in bacteria based on cell shape or as a physical regulator of eukaryotic cell division in yeast^53-55^. Furthermore, fungal pathogens exhibit mechanosensation by adapting their actin cytoskeleton to overcome challenges from the external milieu. This aspect is particularly pertinent in the context of interactions at the host-pathogen interface, wherein mechanical defenses are primarily employed to abrogate pathogenic invasion. However, present understanding of mechanotransduction-guided developmental implications during pathogenesis is limited, as prior efforts have primarily focused on either a pathogen or host-centric perspective, leaving a more holistic understanding elusive. Our transcriptomic results contrasting the distinct growth conditions in 3D physical confinement against liquid cultures demonstrate that soil-like confinement is a major determinant of the fungal gene expression programs regulating metabolism in response to growth conditions. Importantly, these findings strongly advocate that growth under 3D confinement represents a fundamentally different state of fungal growth characteristic of natural soil environments. Hence, we envision the 3D culture platform presented in this study as an opportunity to move beyond the traditional models of liquid-grown axenic cultures and help redefine our understanding of host colonization by fungi in soil-like microenvironments.

Present knowledge of soil-borne fungal diseases and their infection progress is based on datasets derived from interactions occuring in an opaque medium (soil). This also hampers a systematic characterization of specific root zone preference by the invading fungus, which is largely described as being strain dependent. Our platform bridges this gap by quantitatively evaluating progression of vascular wilt infection by *F. oxysporum* and the associated pathogenicity symptoms using *in-planta* infection assays in tomato. Here, we confirm the progression of clear wilting symptoms as well as the evaluation of infection by a fungal isogenic mutant that is compromised in virulence. Moreover, we accomplish both direct visualization and molecular characterization of early infection events leading to colonisation of the tomato cortical cells as well as xylem-mounted host defense responses against fungus in proximity to the plant root. Hence, our 3D platform represents a transformative tool for dissecting the early signalling events in pathogenesis, as well as a tractable approach towards resolving open questions in infection mechanobiology which are otherwise challenging to investigate in natural soil-like setting.

Despite our primary understanding of immune responses triggered in response to plant-fungal interactions being limited to events following physical contact or invasion^56-57^, the recognition and sensing of microbes likely occurs even before physical contact is established. However, quantitative spatial descriptors of these sensing modalities as well as the physical boundaries of such interactions remain poorly defined. Our work here conclusively demonstrates proximal re-orientation of *F. oxysporum* hyphae towards the root tip at distances of ∼500 µm. Our approach also identifies new *ex-planta* subsets of secreted proteins from *F. oxysporum* across different zones of root tissue, in which putatively guide physical contact with and adherence to the plant root. Our findings on the plant defense responses constitute an important conceptual advancement demonstrating that distinct immune signaling responses arise from developmentally distinct root zones, representing a channelized modular pattern instrumental for maintaining a growth-defense tradeoff. These new pathogen/host datasets obtained from 3D mechanical milieus will be a critical resource for identifying putative mechanosensors as well as deciphering mechanistic insights on their roles in distinguishing root-proximal friend/foe entities.

Altogether, our findings reveal the presence of a hitherto unexplored regime of microbial interactions, which presents significant opportunities for fundamental discoveries into the mechanobiology of growth regulation and infection in filamentous pathogens. Our versatile 3D platform offers an unprecedented ringside view of the pathogen infection processes, which allows spatiotemporal correlation of the transcriptomic landscape with direct visualization. We anticipate that our advancement, combining bioengineered materials with molecular techniques, will enable timely investigations on fungal adaptability in response to soil water/nutrient distribution, soil compaction, as well as microbial confrontation due to native microbiota. Together, these efforts will impart major agricultural benefits, by opening avenues to decipher drought and disease-resistance mechanisms, as well as guiding the development of climate-resilient crops. Importantly, such a system will also provide unparalleled opportunities for elucidating the physical and genetic principles enabling pathogen adaptation to trans-kingdom hosts.

## Materials and Methods

### Preparation of jammed agarose microgels

Jammed agarose microgels were prepared by a process of flash solidification^50^, wherein a hot liquid agarose solution is sprayed into a bath of cold water/media while being continuously stirred. This generates a high Reynolds number regime, which breaks up the stream of liquid agarose into micron-sized droplets that solidify instantly due to the large temperature difference. These flash-solidified microparticles can be subsequently concentrated and collected following centrifugation to generate a jammed 3D packing. This mass represents the 3D growth media, which is a granular, porous matrix. Our strategy also enables a rapid and flexible method to alter the liquid phase composition by following a media exchange step. For this, the 3D matrix prepared in a given aqueous liquid (e.g., water) was mixed with twice the volume of a different aqueous solution (e.g., Gamborg medium), vortex mixed to ensure homogeneous dispersion, and subsequently concentrated via centrifugation. This process was repeated twice to ensure effective replacement of the aqueous phase.

### Rheological characterization of 3D growth media

We used an Anton Paar 302e rheometer to characterize the rheological properties of the jammed 3D microgel. All measurements were performed using a roughened cone plate system at room temperature in a shear-controlled mode. First, we evaluated the bulk viscoelastic properties of the matrix by applying a small amplitude (1%) oscillatory shear over different frequencies, while recording the complex shear moduli – this can be resolved into specific components of elastic storage modulus (G’) and viscous loss modulus (G’’). While the G’ represents the energy stored during a deformation cycle and hence corresponds to the elastic properties of the material, the G’’ represents the energy dissipated during deformation and reports on the viscous fluid-like behaviour of the material (Bhattacharjee et al., 2018). It naturally follows that G’ > G’’ suggests an elastic solid-like nature, which is observed using our 3D growth media. Hence, jammed agarose microgels at rest behave as a soft solid. We also observed that both the G’ and G’’ remained largely conserved across different frequencies of shear, indicating that the shear moduli of the system were independent of the shear rate, suggesting that the material has time-invariant mechanical properties^54^.

Since plants and fungi under 3D confinement need to locally deform the surrounding matrix during growth, we also evaluate the material’s shear stress response to different shear rates. For this, we apply a unidirectional shear at varying rates while recording the shear stress response^58^. For a high shear rate, we observe a proportionate change in the stress response, indicating a fluid-like behaviour. However, under low regimes of shear, the stress response plateaus, indicating a soft, solid-like nature. Hence, our jammed 3D microgel can reversibly transition between a liquid-like and a solid-like state^54^. This transition point – the shear stress above which the material flows and below which it remains solid – is referred to as the yield stress of the system, and the shear rate at which this transition occurs is referred to as the crossover shear rate. The yield stress behaviour of the 3D microgel media, hence, allows for localized fluidization followed by immediate recovery^59^, which is critical for both *in situ* plant and fungal growth and routine experimental manipulations without altering the bulk material properties.

### Microparticle morphometry and pore size quantification

For characterizing the morphological attributes of individual micro particles, we prepare a dilute suspension (1:1000, v/v%) of the jammed agarose microgels in distilled water. This is thoroughly vortex mixed to break up any clumps and ensure homogeneous particle dispersion. Following this, we image ∼200 µL of this suspension using a phase contrast-equipped wide field microscope (Nikon TE300 Eclipse). For quantitative measurements, we manually trace the outlines of these particles in ImageJ and obtain values for specific shape descriptors using the in-built particle analysis functions. Given the expected microporous nature of our 3D matrix, we probe the disordered pore space geometry using 200 nm fluorescent tracer particles. A dilute (1:100 final concentration from a 1% w/v stock solution) suspension of these beads is prepared using 200 µL of the jammed agarose microgels in a glass-bottom imaging dish. We acquire high-speed (∼19 fps) time-lapse imaging of the thermal energy-driven diffusion of such beads within the pore spaces of the 3D growth media. For each such bead, we employ custom MATLAB scripts based on the Crocker-Grier algorithm to track the bead centre with sub-pixel accuracy and calculate its displacement. From this data, we obtain the mean square displacement (MSD) over time, and plot this as a function of different time periods. The MSD values increase linearly with time during unimpeded motion until an eventual plateau. Physically, this corresponds to the bead diffusion being impeded by the walls circumscribing each pore. The square root of this plateau MSD value added to the bead diameter directly gives the smallest constraining dimension of the pore space explored by the diffusing bead.

### Plant growth conditions

Tomato seeds (cv. Moneymaker) were sterilized by adding 0.1% bleach solution as previously reported^33^. The sterilized seeds were then transferred to a container with 50ml sterile water, covered with foil and incubated 28°C and shaken at 150 rpm for 3-4 days. For the comparative substrate growth experiment, pre-germinated seeds were either grown in Soilrite mix (Keltech Energies Ltd, Bangalore, India), ½ MS agar (Himedia, PCT0901) without sucrose (Himedia, PT025) or microjammed gel in Gamborg (Duchefa, G0210.0050) in 15ml centrifuge tubes. Upon the emergence of the first pair of true leaves, the plants were uprooted, and the roots were washed and patted dry before further experiments. For experiments involving the pipette set-up, the pre-germinated seedlings were transferred to a petri dish lined with a wet filter paper. Upon emergence of the shoot, the seedlings were then placed in the pasteur pipette bulb and covered with parafilm to maintain humidity. The cover was removed once the cotyledons began to emerge. The pipettes were watered with sterile water every 24 to 48 hours. The seedlings were grown at 28 °C and 70% humidity under a 16:8-hour day-night photoperiod.

### Chlorophyll quantification

Two-week-old tomato seedlings grown in different substrates were submerged in methanol and placed in the dark at 4°C for 24 hrs. The extracts were then measured for their absorbance at 665 nm (chlorophyll a) and 652 nm (chlorophyll b) using a Thermo Scientific Varioskan LUX Multimode Microplate Reader. The total amount of chlorophyll was calculated per plant as described previously^60^.

### DAB staining

Two-week-old tomato seedlings were stained with 3,3’-diaminobenzidine (DAB) (Sigma, D5637). For positive control, roots were treated with 200 nM *flg22* peptide (GenScript, RP19986) for 30 minutes. DAB solution (1 mg/mL) was made by dissolving the powder in ddH_2_O. The roots were then submerged in the solution and were vacuum infiltrated for 30 min. The roots were then incubated at RT for 30 minutes. The roots were destained with ethanol. Roots were then mounted and imaged under a bright field using a Zeiss Axioscan7 microscope.

### Fungal strains, growth conditions and pathogenicity

For visualization experiments, Fol-3xmClover strain was used^61^. For pathogenicity tests, wild-type Fol4287and the Δ*fmk1* mutant in the same background were used. The fungal strains were grown in Potato Dextrose Broth (HiMedia, GM403) by inoculating a stock culture with or without antibiotic selection. The cultures were then incubated for 72 hours at 28°C and shaken at 150 rpm. The spores were pelleted by centrifugation and resuspended in sterile water to achieve a spore inoculum of 10^5^ spores/mL. For infections in the pipette set-up, fungal spores were inoculated in the vicinity of the root. After 35 days, the number of plants in each stage was counted for the disease index using the disease scheme. The disease progression scheme is as follows: stage 1 – plants are green; stage 2 – yellowing in 1-2 leaves; stage 3 – yellowing and wilting in more than 2 leaves; stage 4 – loss of leaves, wilting, stunted growth. For assessing pathogenicity, the fresh weight of the plants was recorded. For microscopy assays and transcriptomic studies, the pre-germinated seedlings were transferred to 35mm glass-bottom dishes with the root tips touching the glass surface and covered with Gamborg-based microgel mixed with spore inoculum of 10^5^ spores/mL with the desired strain.

### Fungal growth curve assays

Fungal spores were mixed in Gamborg liquid and microgel. The samples were then transferred to a 96-well plate (200 µL in each well) with 3 technical replicates for each condition. The liquid samples were incubated at 28°C and shaken at 150 rpm. The microgel samples were incubated at 28°C without shaking. The measurements for fluorescence (488 nm) and absorbance (600 nm) were taken every 24 hrs using the Thermo Scientific Varioskan LUX Multimode Microplate Reader. Analysis and representation of the results were performed using OriginLab software.

### Image acquisition

All microscopy images in this study were acquired using a point-scanning laser confocal microscope (Nikon, A1HD25), barring those for the microparticle morphometry, which were acquired using a phase contrast-equipped widefield microscope (Nikon, Ti2 Eclipse). For image acquisition, all samples are prepared in glass-bottom 35 mm dishes as described previously. Phenotypic images of plants were captured using a Nikon D5700 camera.

### Gene Expression Analysis using RT-qPCR

Plants were uprooted from the microgel setup, and the gel particles were washed off. At every time point, roots from 5 plants were pooled and RNA was extracted from roots using the RNeasy Kit (Qiagen, 74104) with at least 100 mg tissue/gel and 500 µL of lysis buffer. RNA was eluted in 40 µL nuclease free water, followed by DNase treatment. cDNA synthesis was carried out using PrimeScript 1st strand cDNA Synthesis Kit (Takara Biosciences, 6110A). qPCR was performed with the iTaq Universal SYBR Green Supermix (Bio-Rad, 172-5121) and 0.2 µM of forward and reverse primers (Eurofins Genomics, India). The primers used in the study are listed in the SI Data Table 1. The PCR was performed with an initial denaturation at 95°C for 10 minutes, followed by 40 cycles of 95°C for 10 seconds, 60°C for 10 seconds, and 72°C for 20 seconds. Sl*-GAPDH* was used as an internal control for profiling of tomato defense genes. Gene expression was calculated using the ΔΔCt method.

### Library preparation & sequencing

For the spatiotemporal transcriptomics, plant and fungal RNA were isolated from the selected time point and samples using the Qiagen RNeasy kit according to the manufacturer’s instructions. The RNA quantification was performed on a Qubit fluorometer, and integrity was evaluated on an Agilent TapeStation. All treatments were performed for 3 biological replicates. For comparison of Fol growth *in vitro*, in Gamborg liquid and microgel, fungal samples from 3 replicates were harvested for the study. For the plant-fungus interaction study, the fungal samples from 25 individual plants and proximal/distal fungal hyphae surrounding roots from 25 petri dishes were pooled to make one biological replicate and three such biological replicates were used for the experiment. From the total RNA, mRNA was enriched using the NEBNext Poly(A) mRNA magnetic isolation module. The library was prepared using NEBNext Ultra II Directional RNA Library Prep with sample purification beads. The library was sequenced on the NovaSeq 6000 platform using SP flowcell with 2×100 bp paired-end reads with a depth of 30 million.

### Transcriptomic Analysis and Visualization

The raw reads were quasi-mapped against the reference transcriptomes of *Solanum lycopersicum* (SL 4.0) and *F. oxysporum* f. sp. *lycopersici* (GCF_000149955.1) downloaded from Phytozome and NCBI, respectively, using Salmon Quant^62^. The raw reads have been uploaded to the ArrayExpress database. DESeq2^63^, along with the tximport package from RStudio, were used to perform differential gene expression analysis. After pairwise differential gene expression analysis, genes with log2FC >2 or < −2 and p-adj <0.05 were denoted as differentially up- or down-regulated, respectively. When comparing across samples, gene expression values are represented as Z-scores (standard scores), which are calculated by subtracting the mean read count across all samples from the read count in the sample and dividing that by the standard deviation. For the confined state experiment, *Fol* grown in minimal medium shaken at 150rpm was used as a reference for normalizing *Fol* grown in liquid or gel Gamborg medium. The DESeq2 output is listed in **Supplemental Table 1**. The list of fungal stress markers identified in our dataset is mentioned in the **SI Data Table 2**. For the fungus recovered from the roots of infected samples or the microgel, the fungus grown in the microgel without any plant was used as a reference. The DESeq2 output is listed in **Supplemental Table 2**. For plant samples, uninfected root and tip tissues were used as references for infected root and tip tissue samples, respectively. The DESeq2 output is listed in Supplemental Table 3. All the plots have been prepared in RStudio/Python with custom scripts. Gene Ontology enrichment analyses were performed using ShinyGO64 with an FDR cut-off of 0.05. InterPro Term Enrichment was carried out using the func-e package from Python. The entire proteomes were flushed through InterPro (standalone version), and the output was used as background to perform the enrichment in the selected genes. All downstream format conversions and visualizations were either done manually or by custom R/Python scripts. Effectors were predicted by running all the secreted up-regulated genes through SignalP3^65^ to filter for secreted proteins and then EffectorP 3.0^66^ to identify potential effector candidates. The effectors identified in all conditions relative to only fungus control are listed in the **SI Data Table 3**. All scripts utilised in this work have been uploaded to GitHub (https://github.com/ameyred/Microjammed-Gel-For-Fol-Tomato-Interaction).

## Acknowledgements

We also acknowledge the help from Dr. Diya Binoy Joseph at iBRIC-inStem for help on the Zeiss Axioscan7 microscope to capture DAB images. We thank the Central Imaging and Flow Cytometry Facility (CIFF) at NCBS for access to microscopy, as well as the lab-kitchen facilities at NCBS for their help in petri plate preparations and greenhouse facilities for plant growth. We thank Dr. Awadesh Pandit, Lakshminarayanan CP and the Next Generation Sequencing facility staff at NCBS-TIFR for help with NGS related experiments.

## Author Contributions

A.R. and T.B. supervised the study. A.R., T.B., and M.S. conceptualised the study. A.R., T.B., M.S., and S.T., contributed towards the study design. M.S. and T.B. developed the 3D culture technique and M.S. performed all material characterization. S.T. performed the experiments and analyzed data with help from A.R.. M.S. collected the confocal microscopy images and performed all image analyses with help from T.B.. V.S. analysed all the NGS data related to the RNA-Seq analysis and graphical presentation. A.P. and M.S. performed initial optimisations of the plant growth microgel setups. S.T., M.S., and A.P. curated the data. S.T. and M.S. prepared the figures, with help from A.R. and T.B.. A.R., M.S., T.B., and S.T., wrote the original manuscript. A.R. and T.B. acquired primary funding support for the project. All authors participated in reviewing and editing the final version of the manuscript.

## Corresponding Authors

Correspondence to Tapomoy Bhattacharjee and Amey Redkar

