## Supplementary Information for "Dissecting the root-fungal interface in 3D reveals spatially distinct signalling landscapes"

This PDF file includes:
Supplemental Information Figs. **4**
Supplemental Data Tables. **3**

**Supplemental Fig 1.**

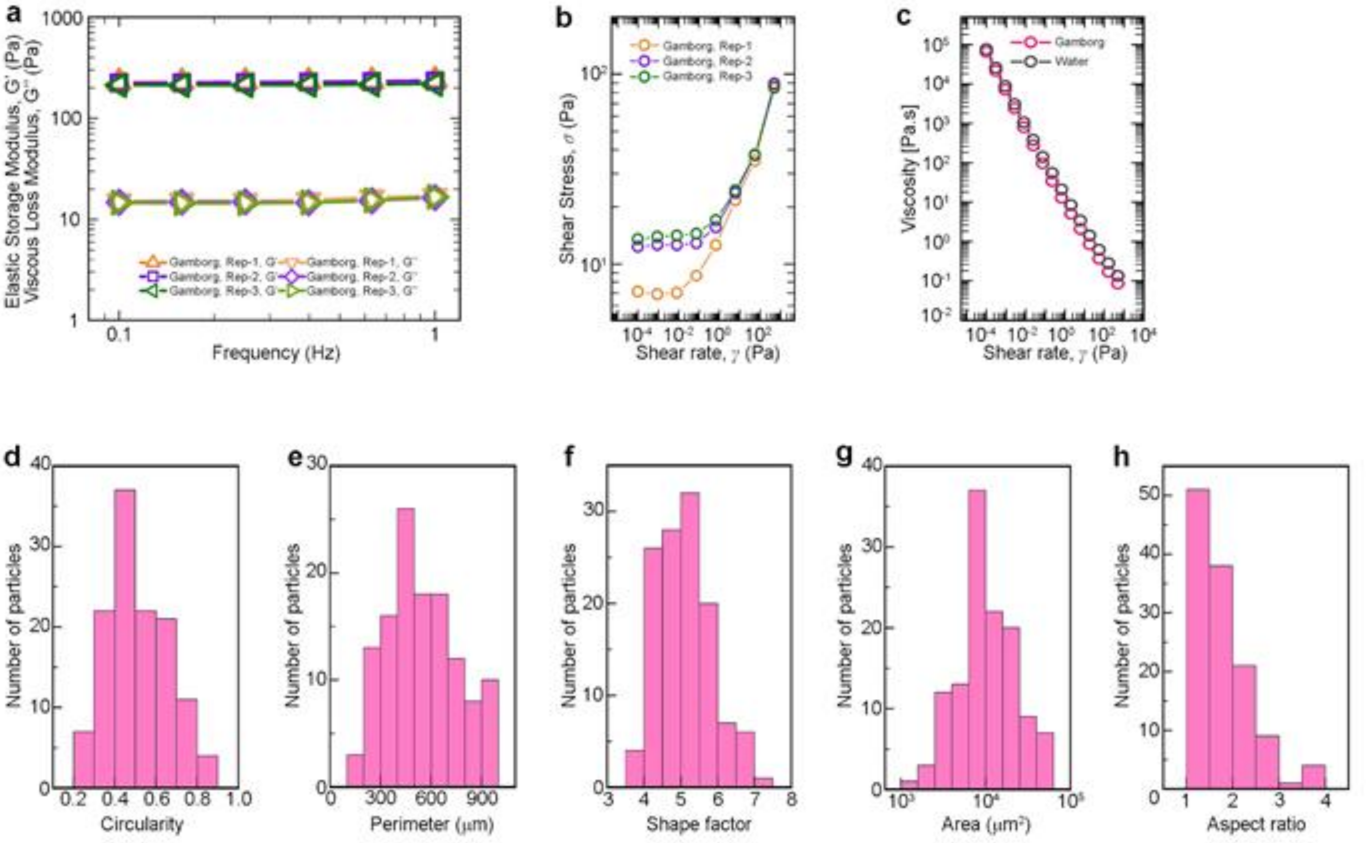


**Supplemental Fig 1 | Rheological characterization of the Gamborg-based 3D microgel. a-b,** Reproducibility of rheological properties demonstrated by independent replicates of Gamborg-infused 3D microgels, synthesized using water as the continuous phase and subsequently subjected to media exchange with Gamborg nutrient broth (see Materials and Methods). **c,** Viscosity of 3D microgels recorded across different shear rates, synthesized using either water or Gamborg media as the continuous phase. **d-g,** Morphological analysis of the microgel particles. (**d)** Circularity, (**e)** Perimeter, (**f)** Shape factor, (**g)** Area, and (**h)** Aspect ratio.

**Supplemental Fig 2.**

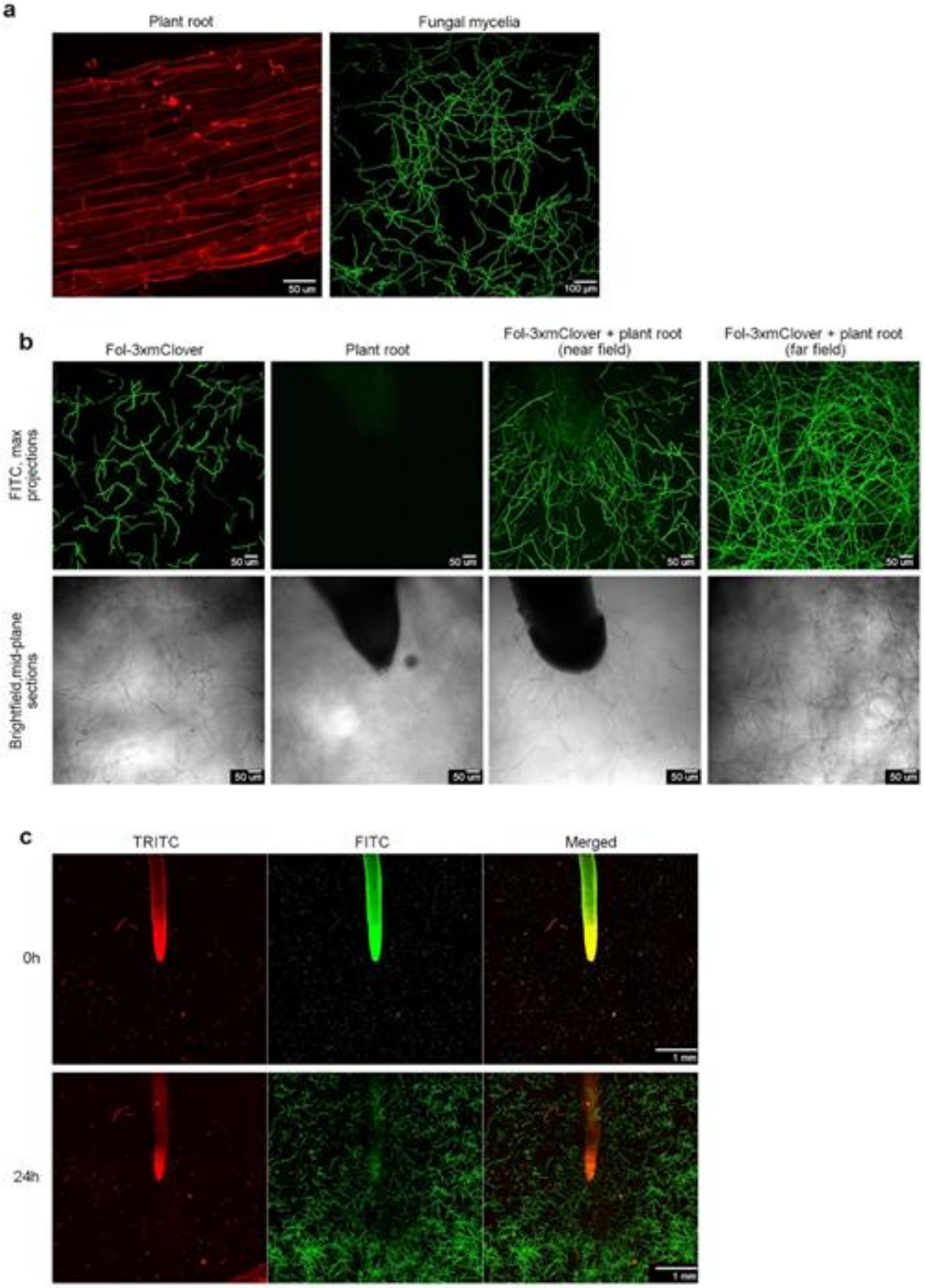


**Supplemental Fig 2 | 3D imaging of *F. oxysporum* growth and development reveals a fungal modulation in response to host roots. a,** Visualization of plant root (stained with propidium iodide) and fungal mycelia (Fol-3xmClover3) grown in Gamborg-based 3D microgel. **b,** Re-orientation of the *F. oxysporum* f.sp. *lycopersici* (Fol)-3xmClover3 hyphae, in presence of tomato root. Plates with Gamborg-based 3D microgel are incubated either with (Fol)-3xmClover3 spores alone or tomato root alone or in presence of both. Near and far field confocal images corresponding to the same plate imaged close to and further away from the root are shown in brightfield and fluorescence channels, respectively. Images are captured at 36 hours post infection (hpi) to highlight the re-orientation of the fungal hyphae, only in the presence of the tomato root. Scale bar = 50µm. **c,** Time-lapse confocal imaging of root–fungus interaction over 24 hours. Root tips (red) are imaged at 0 and 24 hpi with (Fol)-3xmClover3 (green). Images at the 0-hour time-point show the initial state of the tomato root with intact root tips and minimal fungal presence. At the 24-hour time-point, fungal hyphae re-orient around the root surface towards the host root tip.

**Supplemental Fig 3.**

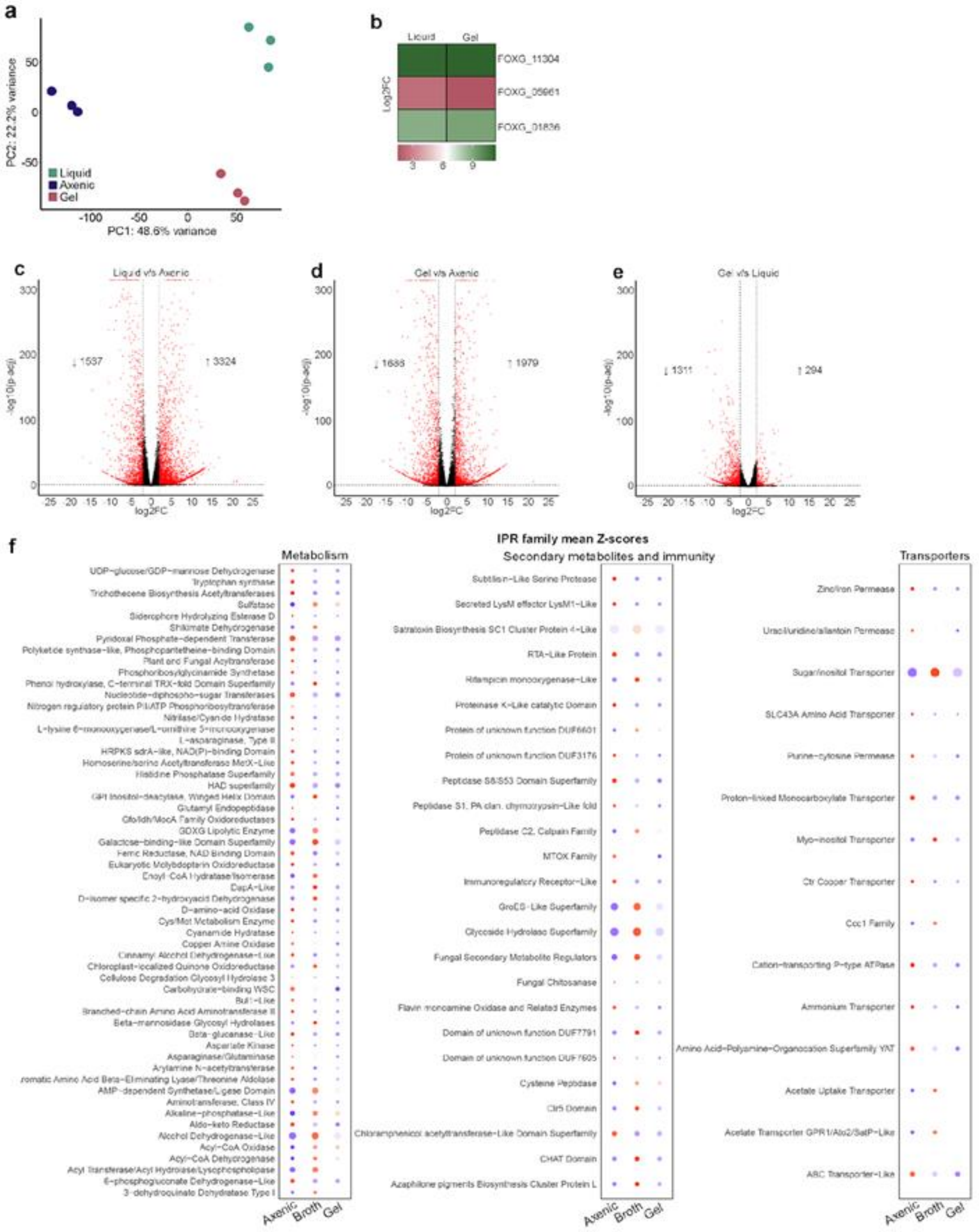

**Supplemental Fig 3 | Transcriptional dynamics in *F. oxysporum* growth and development during liquid and confined state. a,** Principal Component Analysis (PCA) plot showing $Log_2$-transformed read counts and the transcriptional reprogramming of gene expression between axenic, liquid and microgel. Axenic represents *F. oxysporum* f.sp. *lycopersici* (Fol) grown in liquid minimal medium. Liquid represents Fol grown in liquid Gamborg B5 medium. Microgel represents Fol grown in microgel jammed Gamborg B5 medium. **b,** $Log_2$-transformed fold changes in transcript levels of Fol genes encoding for stress markers from a list of 78 genes identified across fungi. Note the absence of differences in their expression level across the samples tested, indicating a stress-free growth. **c-e,** Volcano plots showing pairwise differential expression analysis of Fol genes after $DESeq_2$ under different growth conditions at 24 hours. Microgel v/s axenic represents Fol grown in Gamborg microgel, normalised to Fol grown in liquid minimal medium. Liquid v/s axenic represents Fol grown in liquid Gamborg medium, normalised to Fol grown in liquid minimal medium. Microgel vs. liquid represents Fol grown in Gamborg microgel, normalised to Fol grown in liquid Gamborg medium. All liquid cultures are shaken at 150 rpm. **f,** DEGs across growth conditions represented as the mean Z-score for each IPR family across conditions. The IPR terms showed an enrichment of genes classified into transporters, metabolism, secondary metabolites and immunity-related genes.

**Supplemental Fig 4.**

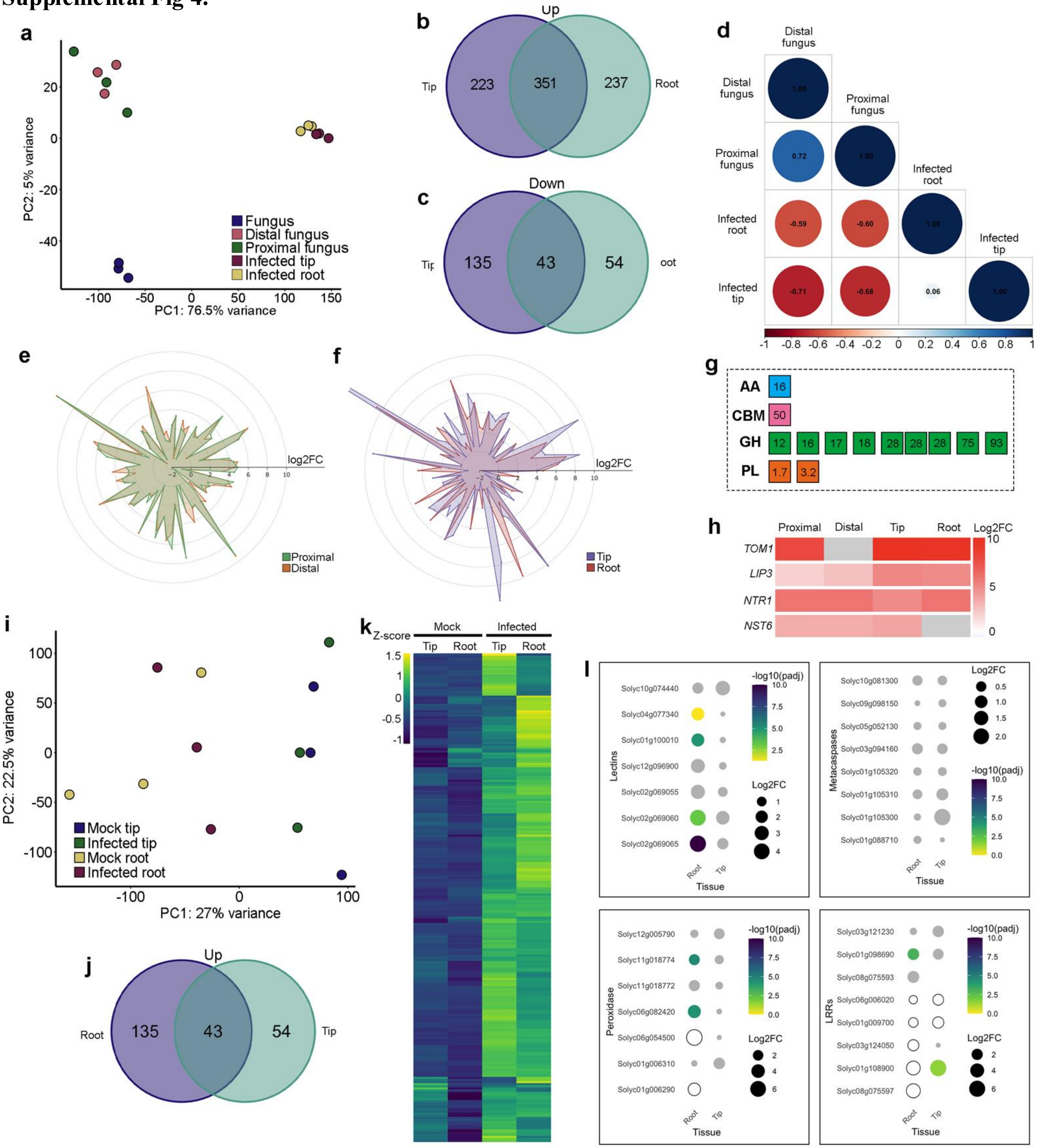

**Supplemental Fig 4 | Host and pathogen early signalling landscape of *F. oxysporum* infected tomato showing *ex-* and *in-planta* signalling modules a,** Principal Component Analysis (PCA) plot showing $Log_2$ transformed read counts and the transcriptional reprogramming of gene expression between only fungus, proximal fungus, distal fungus, fungus in infected root tip and bulk samples. Each circle represents a biological replicate, and samples are coloured by treatment. **b-c**, Venn diagram of Fol genes encoding predicted secreted proteins (SPs) up-regulated **(b)** and down-regulated **(c)** in tomato tip (purple) vs. bulk of the roots (green) at 24 hours post inoculation (hpi) versus axenic culture of only fungus in Gamborg microgel. **d,** Correlation matrix plot showing the Pearson's correlation coefficient of all the identified secreted proteins between Fol samples from different fungal regions *ex-planta* (proximal and distal) in relation to the plant root (tip and bulk root). **e-f,** Radar plots representing the overlap between Fol secreted protein genes up-regulated *ex-planta* near the root (proximal) and as compared to the fungus growing away from the root (distal) **(e)** and Fol secreted protein genes up-regulated *in-planta* within the root tip and the root bulk. Each axis represents a single gene plotted with log2FC values. **g,** Candidate Fol effectors identified in *ex-* and *in-planta* dataset belonging to CAZyme families. Each box represents one effector, and the number indicates the family; AA-Auxiliary Activities, CBM-Carbohydrate-binding module, GH-Glycoside hydrolase, PL-Polysaccharide lyase. **h,** PCA plot showing $Log_2$ transformed read counts and the transcriptional reprogramming of gene expression between mock and infected root and mock and infected tip samples. Each circle represents a biological replicate, and samples are coloured by treatment. **i**, Venn diagram of tomato genes unregulated genes in the infected root tip (green) versus the root bulk (purple) at 24 hours post inoculation versus the non-infected mock control. **j,** Heat map of the DEGs in tomato as a comparison of mock v/s infected root samples as plotted using their mean z-scores. **k,** Gene expression patterns of lectins, metacaspases, peroxidases and leucine-rich-repeats (LRRs) across root tip and bulk root samples. Gene expression is represented as $Log_2FC$ values. Genes with p-adj > 0.05 marked as non-significant (ns) and represented as grey. Genes marked in white do not have any padj value. Absence of a dot indicates that $Log_2FC$ value for the gene was not detected in that sample.

**Supplemental Data Tables**

**Supplemental Data Table 1. Primers used in this study.**

| Gene | Gene ID | Direction | Sequence | Amplicon size | Source |
|---|---|---|---|---|---|
| *GAPDH* | Solyc05g014470.2 | Forward | GGCTGCAATCAAGGAGGAA | 204 | Expósito-Rodríguez, *et al.,* (2008) |
| | | Reverse | AAATCAATCACACGGGAACTG | | |
| *PR1* | Solyc09g007010.1 | Forward | TCTTGTGAGGCCCAAAATTC | 245 | Aime et al., (2013) |
| | | Reverse | TAGTCTGGCCTCTCGGACA | | |
| *CHI3* | Solyc02g082920.2 | Forward | TGCAGGAACATTCACTGGAG | 248 | Aime et al., (2013) |
| | | Reverse | TAACGTTGTGGCATGATGGT | | |
| *GLUA* | Solyc01g008620.2 | Forward | GTCTCAACCGCGACATATT | 249 | Aime et al., (2013) |
| | | Reverse | CACAAGGGCATCGAAAAGAT | | |

**Supplemental Data Table 2. Fungal stress markers.**

| Stress Markers | Liquid_L2FC | Liquid_padj | Gel_L2FC | Gel_padj |
|---|---|---|---|---|
| FOXG_01684 | #N/A | #N/A | #N/A | #N/A |
| FOXG_14703 | #N/A | #N/A | #N/A | #N/A |
| FOXG_15645 | #N/A | #N/A | #N/A | #N/A |
| FOXG_11304 | 11.15140094 | 4.47145E-20 | 11.29268987 | 1.81448E-20 |
| FOXG_01836 | 8.849895093 | 3.4695E-106 | 9.175167021 | 1.11404E-38 |
| FOXG_05961 | 2.277436385 | 7.44606E-23 | 1.465227201 | 1.76183E-28 |
| FOXG_02214 | 1.77929182 | 7.18236E-06 | 1.169776512 | 0.024360902 |
| FOXG_11273 | 1.013258896 | 3.24885E-16 | 0.518170934 | 1.36886E-05 |
| FOXG_01993 | 0.832353229 | 2.27838E-11 | 0.322363148 | 0.000447531 |
| FOXG_01632 | 0.795545736 | 3.50382E-11 | 0.243352281 | 0.02050388 |
| FOXG_02420 | 0.775309442 | 0.002879548 | 0.839056094 | 0.00265798 |
| FOXG_06235 | 0.674646035 | 7.06444E-05 | 0.803433728 | 4.78232E-06 |
| FOXG_05428 | 0.57460752 | 6.06971E-09 | 0.432059186 | 1.25439E-06 |
| FOXG_00281 | 0.494855885 | 0.000886716 | 0.506238501 | 0.000330692 |
| FOXG_01684 | 0.487862869 | 1.52205E-07 | -0.125214273 | 0.323999119 |
| FOXG_06022 | 0.451622435 | 0.002466472 | 0.32343753 | 0.141250252 |
| FOXG_06370 | 0.398237086 | 0.001758359 | 0.274964352 | 0.110436592 |
| FOXG_08078 | 0.39685947 | 0.000239054 | 0.312794615 | 0.005579282 |
| FOXG_15663 | 0.360173832 | 8.98162E-05 | 0.490106955 | 8.33738E-05 |
| FOXG_06120 | 0.350244887 | 8.68184E-07 | 0.191906896 | 0.016359131 |
| FOXG_18412 | 0.292041543 | 0.000319764 | 0.062416572 | 0.559498472 |
| FOXG_05887 | 0.278954285 | 0.019059045 | 0.189179423 | 0.565427811 |
| FOXG_05278 | 0.277992721 | 0.018673104 | -0.270543015 | 0.026505254 |
| FOXG_07912 | 0.222837771 | 0.142267093 | -0.25582425 | 0.054697887 |
| FOXG_03010 | 0.218376511 | 0.177836119 | 0.283460594 | 0.089454884 |
| FOXG_00016 | 0.206396159 | 0.072099214 | 0.116816092 | 0.351219461 |
| FOXG_04163 | 0.203221164 | 0.040432958 | -0.06042777 | 0.527235295 |
| FOXG_03343 | 0.154630389 | 0.054943509 | -0.103488478 | 0.483218784 |
| FOXG_02117 | 0.129387939 | 0.179148848 | -0.10765612 | 0.343798478 |
| FOXG_11105 | 0.113297316 | 0.337803317 | 0.12172842 | 0.189598702 |
| FOXG_13835 | 0.084169566 | 0.213915829 | 0.425242331 | 4.23964E-11 |
| FOXG_01859 | 0.084038835 | 0.506995842 | 0.071320275 | 0.48706925 |
| FOXG_01907 | 0.081293326 | 0.535318314 | -0.224023328 | 0.024984688 |
| FOXG_09534 | 0.058374413 | 0.611954125 | 0.133855168 | 0.062044209 |
| FOXG_03067 | 0.05784047 | 0.643360694 | -0.462939682 | 0.000221892 |
| FOXG_08113 | 0.052924093 | 0.809629241 | -0.327805651 | 0.073340899 |
| FOXG_10511 | 0.009349285 | 0.947779718 | 0.044149317 | 0.740759903 |
| FOXG_11532 | -0.021810462 | 0.818133967 | -0.005673894 | 0.96893411 |
| FOXG_05092 | -0.039689258 | 0.681840935 | -0.227634099 | 0.050529144 |
| FOXG_08584 | -0.117723786 | 0.344343995 | 0.006580908 | 0.972653651 |
| FOXG_07983 | -0.129972043 | 0.774029611 | -0.346399008 | 0.330046442 |
| FOXG_06419 | -0.181221587 | 0.286890861 | 0.73662979 | 1.68732E-07 |
| FOXG_02319 | -0.198654049 | 0.097086065 | 0.013469197 | 0.949181484 |

| | | | | |
|---|---|---|---|---|
| FOXG_01955 | -0.206604653 | 0.047139069 | -0.252877599 | 0.079244072 |
| FOXG_04162 | -0.210534077 | 0.150419288 | -0.396460264 | 0.000222389 |
| FOXG_09254 | -0.239988828 | 0.001668731 | -0.205226426 | 0.03245171 |
| FOXG_01180 | -0.258374323 | 0.011649233 | -0.601126972 | 3.35231E-07 |
| FOXG_00578 | -0.322055318 | 0.014052838 | -0.282128333 | 0.016799963 |
| FOXG_12436 | -0.340820326 | 0.555377661 | -0.448963418 | 0.460079377 |
| FOXG_11409 | -0.385495403 | 0.005880435 | -0.651423816 | 9.53076E-09 |
| FOXG_02224 | -0.418580773 | 0.000423237 | -1.69441497 | 2.97841E-62 |
| FOXG_08140 | -0.476397724 | 4.49573E-05 | -0.592550735 | 1.41632E-08 |
| FOXG_05408 | -0.478493637 | 2.01583E-05 | -0.532646937 | 1.59099E-05 |
| FOXG_02952 | -0.534002567 | 0.489806617 | -0.454327068 | 0.45964356 |
| FOXG_10529 | -0.535267809 | 5.43567E-05 | -0.174606281 | 0.02931835 |
| FOXG_01740 | -0.556584857 | 0.000293579 | -1.423621487 | 8.2703E-42 |
| FOXG_03261 | -0.656184234 | 0.00443178 | -0.544718753 | 6.83698E-06 |
| FOXG_07703 | -0.779899304 | 2.05818E-13 | -0.252648276 | 0.011374209 |
| FOXG_00919 | -0.98371025 | 9.22106E-28 | -0.729414032 | 0.000681929 |
| FOXG_05265 | -0.993332656 | 3.46657E-24 | -2.24802172 | 3.0111E-173 |
| FOXG_07613 | -1.005269858 | 1.77888E-20 | -0.563448568 | 0.002313558 |
| FOXG_13733 | -1.037665593 | 7.66713E-16 | -1.254998627 | 4.58905E-24 |
| FOXG_10351 | -1.077865447 | 3.86484E-23 | -0.790895181 | 2.30878E-13 |
| FOXG_00005 | -1.122156617 | 2.84527E-17 | -0.770491865 | 6.27261E-11 |
| FOXG_08063 | -1.225887537 | 9.22741E-32 | -0.832121844 | 5.81998E-08 |
| FOXG_01835 | -1.243119455 | 9.79304E-54 | -0.750377418 | 1.56762E-10 |
| FOXG_11503 | -1.411177087 | 5.74189E-28 | -0.562758894 | 3.09873E-11 |
| FOXG_05886 | -1.436223415 | 4.6314E-53 | -1.480459919 | 2.55486E-59 |
| FOXG_06318 | -1.669997803 | 1.6256E-124 | -1.792200253 | 3.06562E-54 |
| FOXG_18886 | -1.980504698 | 2.45619E-35 | -2.838752617 | 1.00787E-28 |
| FOXG_00394 | -2.066744703 | 9.39563E-36 | -1.413483437 | 2.55558E-45 |
| FOXG_02222 | -2.162959188 | 1.95362E-71 | -2.838898709 | 1.30649E-89 |
| FOXG_07915 | -2.778411614 | 3.62282E-66 | -3.171357418 | 4.43364E-74 |
| FOXG_10353 | -2.88185338 | 1.314E-260 | -1.958306231 | 2.6476E-36 |
| FOXG_10530 | -3.052976917 | 1.60551E-62 | -2.662987128 | 8.62618E-41 |
| FOXG_10428 | -5.196376492 | 9.6464E-242 | -5.295310942 | 1.09884E-92 |
| FOXG_16833 | -5.511970415 | 4.277E-107 | -4.102959408 | 3.27645E-89 |
| FOXG_08636 | -5.715733565 | 6.85524E-07 | -7.434905829 | 1.47296E-08 |

**Supplemental Data Table 3. Effectors identified across conditions.**

| Distal_Fungus | Proximal_Fungus | Infected_Tip | Infected_Root |
|---|---|---|---|
| FOXG_09689 | FOXG_10383 | FOXG_02490 | FOXG_02490 |
| FOXG_02756 | FOXG_12292 | FOXG_12276 | FOXG_12276 |
| FOXG_03723 | FOXG_10144 | FOXG_05861 | FOXG_05861 |
| FOXG_08769 | FOXG_04749 | FOXG_16906 | FOXG_16906 |
| FOXG_12212 | FOXG_09689 | FOXG_02457 | FOXG_02457 |
| FOXG_02457 | FOXG_04115 | FOXG_04988 | FOXG_04988 |
| FOXG_04115 | FOXG_14121 | FOXG_07527 | FOXG_07527 |
| FOXG_10490 | FOXG_14109 | FOXG_11848 | FOXG_11848 |
| FOXG_11848 | FOXG_02829 | FOXG_01130 | FOXG_01130 |
| FOXG_15038 | FOXG_10490 | FOXG_12331 | FOXG_12331 |
| FOXG_01130 | FOXG_04089 | FOXG_02756 | FOXG_02756 |
| FOXG_04089 | FOXG_13519 | FOXG_08769 | FOXG_08769 |
| FOXG_10138 | FOXG_12212 | FOXG_12716 | FOXG_12716 |
| FOXG_10144 | FOXG_04055 | FOXG_05607 | FOXG_05607 |
| FOXG_11642 | FOXG_11697 | FOXG_16569 | FOXG_16569 |
| FOXG_12331 | FOXG_17006 | FOXG_05686 | FOXG_05686 |
| FOXG_14607 | FOXG_13198 | FOXG_09814 | FOXG_09814 |
| FOXG_16906 | FOXG_15038 | FOXG_00666 | FOXG_00666 |
| FOXG_05878 | FOXG_22828 | FOXG_20611 | FOXG_20611 |
| FOXG_05945 | FOXG_22929 | FOXG_04805 | FOXG_04805 |
| FOXG_11697 | FOXG_05878 | FOXG_02802 | FOXG_02802 |
| FOXG_12716 | FOXG_13233 | FOXG_04415 | FOXG_04415 |
| FOXG_13204 | FOXG_10138 | FOXG_05948 | FOXG_05948 |
| FOXG_16569 | FOXG_16906 | FOXG_10728 | FOXG_10728 |
| FOXG_10728 | FOXG_14607 | FOXG_12428 | FOXG_10300 |
| FOXG_13198 | FOXG_11881 | FOXG_12416 | FOXG_03723 |
| FOXG_21878 | FOXG_02802 | FOXG_09638 | FOXG_22884 |
| FOXG_05607 | FOXG_12716 | FOXG_13051 | FOXG_03298 |
| FOXG_22828 | FOXG_08769 | FOXG_11400 | FOXG_05945 |
| FOXG_05686 | FOXG_10728 | FOXG_10034 | FOXG_09982 |
| FOXG_09814 | FOXG_00666 | FOXG_14695 | FOXG_08688 |
| FOXG_10383 | FOXG_01130 | FOXG_08922 | FOXG_11033 |
| FOXG_12292 | FOXG_02457 | FOXG_08899 | FOXG_11881 |

| | | | |
|---|---|---|---|
| FOXG_19080 | FOXG_12331 | FOXG_18684 | FOXG_10750 |
| FOXG_00666 | FOXG_08899 | FOXG_16783 | FOXG_10138 |
| FOXG_13519 | FOXG_03723 | FOXG_14607 | FOXG_15793 |
| FOXG_08899 | FOXG_09814 | FOXG_10052 | |
| FOXG_12276 | FOXG_11033 | FOXG_12372 | |
| FOXG_17194 | FOXG_16569 | | |
| FOXG_20611 | FOXG_11400 | | |
| FOXG_22047 | FOXG_16783 | | |
| FOXG_11033 | FOXG_02756 | | |
| FOXG_02829 | FOXG_12416 | | |
| FOXG_11044 | FOXG_05945 | | |
| FOXG_13233 | FOXG_20611 | | |
| FOXG_02802 | FOXG_12276 | | |
| FOXG_11881 | FOXG_11848 | | |
| FOXG_04415 | FOXG_05686 | | |
| FOXG_20247 | | | |
